# Store Independent Ca^2+^ Entry Regulates the DNA Damage Response in Breast Cancer Cells

**DOI:** 10.1101/2020.04.06.027946

**Authors:** Monish Ram Makena, Myungjun Ko, Allatah X. Mekile, Donna K. Dang, John Warrington, Phillip Buckhaults, C. Conover Talbot, Rajini Rao

**Author notes:** Correspondence: Rajini Rao, Ph.D., Department of Physiology, The Johns Hopkins University School of Medicine, 725 N. Wolfe Street, Baltimore, MD 21205. **Abbreviations:** ER, estrogen receptor; PR, progesterone receptor; ROS, reactive oxygen species; SICE, store independent Ca^2+^ entry; SOCE, store operated Ca^2+^ entry; TNBC, triple negative breast cancer; TUNEL, Terminal deoxynucleotidyl transferase dUTP nick end labeling.

## Abstract

Although the mainstay of treatment for hormone responsive breast tumors is targeted endocrine therapy, many patients develop de novo or acquired resistance and are treated with chemotherapeutic drugs. The vast majority (80%) of estrogen receptor positive tumors also express wild type p53 protein that is a major determinant of the DNA damage response. Tumors that are ER+ and p53^WT^ respond poorly to chemotherapy, although the underlying mechanisms are not completely understood. We describe a novel link between store independent Ca^2+^ entry (SICE) and resistance to DNA damaging drugs, mediated by the secretory pathway Ca^2+^-ATPase, SPCA2. In luminal ER+/PR+ breast cancer subtypes, SPCA2 levels are high and correlate with poor survival prognosis. Independent of ion pump activity, SPCA2 elevates baseline Ca^2+^ levels through SICE and drives cell proliferation. Attenuation of SPCA2 or depletion of extracellular Ca^2+^ increased mitochondrial ROS production, DNA damage and activation of the ATM/ATR-p53 axis leading to G0/G1 phase cell cycle arrest and apoptosis. Consistent with these findings, SPCA2 knockdown confers chemosensitivity to DNA damaging agents including doxorubicin, cisplatin and ionizing radiation. We conclude that elevated SPCA2 expression in ER+ p53^WT^ breast tumors drives pro-survival and chemotherapy resistance by suppressing the DNA damage response. Drugs that target store-independent Ca^2+^ entry pathways may have therapeutic potential in treating receptor positive breast cancer.

## INTRODUCTION

Breast cancer is the second most common cancer worldwide. Women born today have a probability of 1 in 8 (12.5%) of developing breast cancer during their lifetime^1^. The majority (70%) of breast cancer is positive for estrogen receptor (ER) and/or progesterone receptor (PR) and is treated with endocrine modulators, including tamoxifen and aromatase inhibitors^2^. Unfortunately, many patients show intrinsic or acquired resistance to hormone therapy or develop serious acute and chronic complications, requiring radiation and chemotherapy^2–4^. There is an urgent need to improve patient outcomes by identifying new biomarker based targets and understanding mechanisms of resistance against current chemotherapeutic agents. In this study we describe a novel role for Ca^2+^ signaling in maintaining genomic integrity to confer pro-survival and chemotherapy resistance in hormone receptor positive breast cancer.

Ca^2+^ is a versatile second messenger that impacts all aspects of cell fate and function^5, 6^. Ca^2+^ homeostasis is maintained by an array of membrane-embedded ion channels, ATP-driven pumps, and carrier proteins that rapidly move Ca^2+^ across organelle or cell membranes. Dysregulation of the expression or activity of Ca^2+^ transporters can produce abnormal elevation or depletion of cellular Ca^2+^ levels, driving oncogenic gene expression, sustaining tumor proliferation and metastasis, and conferring chemoresistance and evasion of apoptosis^2, 7–9^. The presence of microcalcifications, which are radiographically dense deposits of calcium hydroxyapatite crystals in the soft tissue of breast, is an early sign of Ca^2+^ dysregulation and potential malignancy in mammograms^10, 11^.

Central to Ca^2+^ homeostasis mechanisms in breast is SPCA2 (gene name *ATP2C2*), the secretory pathway Ca^2+^ and Mn^2+^ transporting ATPase Isoform 2, found in the Golgi and post-Golgi vesicles of luminal mammary epithelium. SPCA2 expression increases in pregnancy, peaking during parturition, continues to be elevated during lactation for the transport and secretion of Ca^2+^ into milk and returns to basal levels following involution of the mammary gland^12, 13^. In addition to the ATPase-dependent activity required for sequestering Ca^2+^ and Mn^2+^ ions into Golgi and vesicular stores, SPCA2 moonlights as a chaperone and activator of the Orai1 channel through interactions of its non-catalytic N- and C-terminal domains, to elicit robust Ca^2+^ influx at the plasma membrane^14–16^. This mechanism is termed store independent Ca^2+^ entry (SICE) to distinguish it from store operated Ca^2+^ entry (SOCE) that occurs only upon depletion of endoplasmic reticulum Ca^2+^ stores^17^. The coordinated expression of Ca^2+^ transporters and activation of basal to apical Ca^2+^ flux in the mammary gland epithelium is physiologically important for the accumulation of calcium in milk^12^. However, constitutively high expression of SPCA2 was shown to promote tumor growth in receptor positive breast cancers through SICE and downstream Ca^2+^ activation of the MAPK pathway. Thus, silencing of SPCA2 expression in ER+/PR+ xenograft models attenuated tumor formation *in vitro* and *in vivo* ^8, 14^

In this study, we describe a new role for SICE in the DNA damage response pathway. We show that attenuation of SPCA2, or brief depletion of extracellular Ca^2+^, in ER+/PR+ breast cancer cells significantly increased single and double stranded DNA breaks, and activated the ATM/ATR-p53 DNA damage response pathway by eliciting mitochondrial reactive oxygen species (ROS). Consequently, SPCA2-mediated SICE modulates cancer cell resistance to DNA damaging agents, including carboplatin, doxorubicin and ionizing radiation. We uncover an emerging link between store independent Ca^2+^ entry and mitochondrial function that may be critical in blocking ROS-mediated DNA damage. These novel oncogenic functions of SPCA2 are required for cell cycle progression, evasion of apoptosis, and resistance to DNA damaging agents.

## RESULTS

### SPCA2 drives cell cycle progression and survival

Expression of SPCA2 is greatly elevated in the majority of breast cancer subtypes, including ER+, PR+ and HER2+ breast cancer cell lines and tumors^2, 14, 18^, where it confers poor patient survival prognosis (**Fig. S1A**). In contrast, expression of the housekeeping isoform SPCA1 does not alter survival in the same patient population (**Fig. S1B**). Previously, we showed that knockdown (KD) of SPCA2 in the ER+/PR+ breast cancer cell line MCF-7 resulted in reduction in Cyclin D protein, which controls G1 phase progression^14^. Altered expression or activity of cell cycle related proteins and evasion of apoptosis allow cancer cells to divide rapidly and survive^19^. Here, we show that SPCA2 KD significantly reduced cell proliferation in MCF-7 (**Fig. 1A**, **Fig. S1C**). Analysis of the cell cycle revealed a significant increase in the number of cells in the G0/G1 phase from about 40% in control to approximately 60% in the SPCA2 KD cells (**Fig. 1B-D**). Concomitantly, the number of cells that progressed to the S and G2/M phases respectively in SPCA2 KD cells significantly decreased (**Fig. 1B-D**). The ability of SPCA2 to drive proliferation was confirmed in multiple cell lines and was independent of hormone receptor or tumor status: SPCA2 KD inhibited cell growth in the receptor negative, non-cancerous MCF10-A cell line (**Fig. S1D-E**), and in the receptor positive ZR-75 breast cancer line (**Fig. S1F-G**). Furthermore, ectopic expression of SPCA2 in TNBC cell line Hs578T with low endogenous SPCA2^10 18^, greatly increased cell proliferation (**Fig. S1H**).

**Figure 1:**
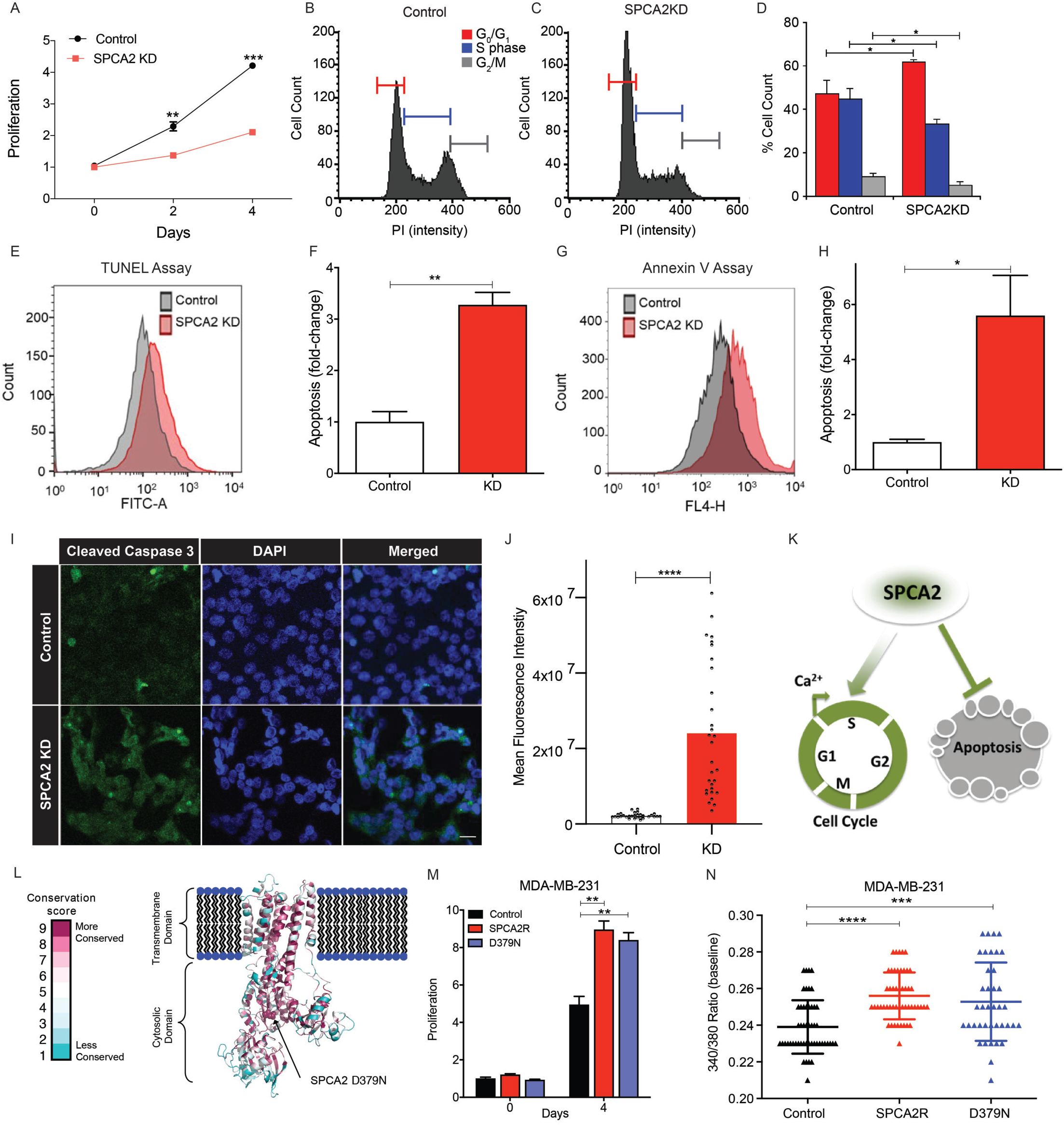
SPCA2 drives cell cycle progression and survival. (A) Cell proliferation in MCF-7 is significantly decreased in SPCA2 KD compared to control, n=3. (B-C) Representative cell cycle distribution (measured by propidium iodide staining) from flow cytometry of control and SPCA2 KD in MCF-7 cells. (D) SPCA2 KD increased percentage of cells in G_o_/G_1_ and correspondingly decreased S and G_2_/M phase cells compared to control, n=3. (E) Distribution of cell count from representative flow cytometry showing apoptosis (measured by TUNEL staining) of control and SPCA2 KD in MCF-7 cells. (F) SPCA2 KD cells significantly increased apoptosis compared to control, n=3. (G) Distribution of cell count from representative flow cytometry showing apoptosis (measured by Annexin V fluorescence) of control and SPCA2 KD in MCF-7 cells. (H) SPCA2 KD cells significantly increased apoptosis compared to control, n=3. (I) Representative confocal microscope images showing immunofluorescence staining of cleaved caspase-3 (marker for apoptosis) in MCF-7 control and SPCA2 KD cells (40x magnification; scale bar, 50 μm). (J) Fluorescence intensity of cleaved caspase-3 labeling was quantified by ImageJ software. SPCA2 KD showed significant increase in cleaved caspase-3 compared to control, n=30 cells. (K) Schematic summarizing data showing that SPCA2 promotes cell cycle proliferation and blocks apoptosis. (L) Schematic of SPCA2 loss of function mutation D379N, located in the highly conserved phosphorylation domain essential for transport activity in all P-type ATPases (M) Overexpression of SPCA2 and SPCA2 D379N in MDA-MB-231 increased cell proliferation compared to control, n=3. (N) Baseline Ca^2+^ levels (340/380 nm ratio of Fura-2 fluorescence) is significantly increased in cells overexpressing SPCA2 and SPCA2 D379N in MDA-MD-231 compared to vector control. Control n=51 cells, SPCA2R n=50 cells, SPCA2 D379N n=39. Significance: *\*P* < 0.05, ** *P* < 0.01, *** *P* < 0.001

One outcome of cell cycle arrest is apoptosis^19^. Hallmarks of apoptosis appeared in MCF-7 following knockdown of SPCA2, as evidenced by increased TUNEL staining (**Fig. 1E-F**), externalization of phosphatidylserine (**Fig. 1G-H**) and staining for cleaved caspase-3 (**Fig. 1I-J**). We conclude that SPCA2 plays a pro-survival role in breast cancer cells (**Fig. 1K**).

The ability of SPCA2 to drive cell proliferation could be due to accumulation of Ca^2+^ in secretory pathway stores. However, ectopic expression of the closely related housekeeping Ca^2+^-ATPase isoform, SPCA1, had no effect on cell proliferation (**Fig. S1I-J**). The isoformspecificity of these observations suggest that luminal Golgi Ca^2+^ is not critical for proliferation, and that SPCA2 may be driving cell growth by a unique, pump-independent mechanism. To test this possibility, we used the previously characterized loss of function mutation D379N^10, 14^, located in the highly conserved phosphorylation domain essential for transport activity in all P-type ATPases **(Fig. 1L**). Expression of mutant D379N in MDA-MB-231 cells which have low endogenous SPCA2 levels^18^ (**Fig. S1K**) increased cell proliferation, similar to wild type SPCA2 (**Fig. 1M**). Previously, we showed that SPCA2 promotes store-independent Ca^2+^ entry through a pumping-independent mechanism. Here, we show that mutant D379N elevates basal Ca^2+^ levels (**Fig. 1N**) and can elicit constitutive Ca^2+^ entry (**Fig. S1L**), similar to wild type SPCA2^18^. Thus, the ability of SPCA2 to elicit robust Ca^2+^ entry and increase baseline cytoplasmic Ca^2+^ levels, independent of ion pumping activity, may drive cell proliferation.

We note that G1 arrest has also been observed upon silencing of STIM1 or Orai1, mediators of store-dependent Ca^2+^ entry (SOCE), in cancer cells^20, 21^. Furthermore, depletion of external Ca^2+^ using the Ca^2+^ chelator EGTA also resulted in reduction of the G1 phase Cyclin D protein^14^. Taken together, these data reveal a critical role for both store-dependent and -independent modes of Ca^2+^ entry in cell cycle progression at the G1/S checkpoint.

**Figure S1:**
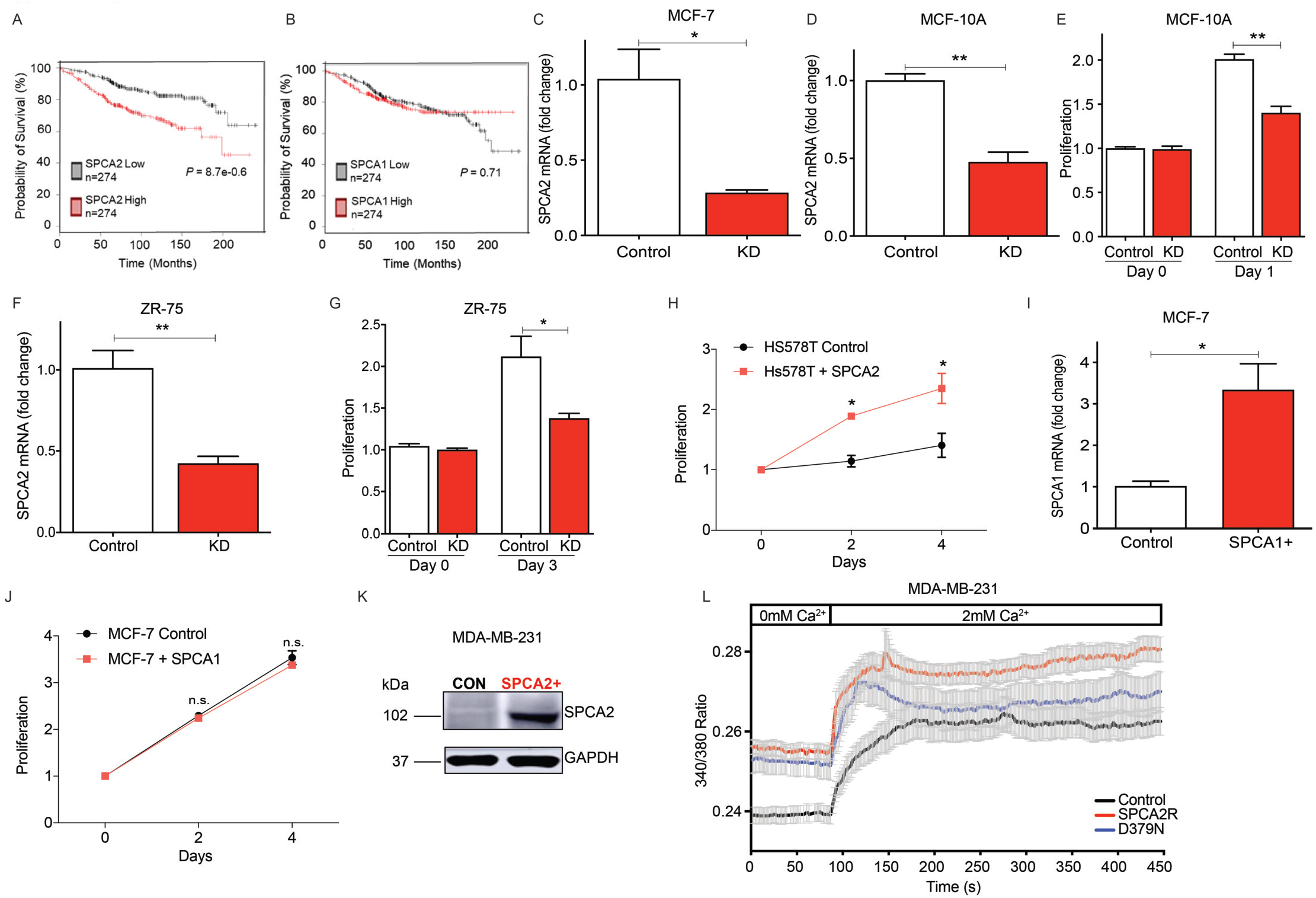
SPCA2 drives cell cycle progression and survival. (A) Elevated SPCA2 is associated with poor overall survival in receptor positive breast cancer subtypes. Data from KMplotter keeping ER+, selected PR (all), and HER 2 (all), and cut off as median. (*P* = 8.7e^−0.6^). (B) Elevated SPCA1 is not associated with poor overall survival in receptor positive breast cancer subtypes. Data from KMplotter keeping ER+, selected PR (all), and HER 2 (all), and cut off as median. (*P* = 0.71). (C) Knockdown of SPCA2 in MCF-7 cells was confirmed by qPCR, n=3. (D) Knockdown of SPCA2 in MCF-10A cells was confirmed by qPCR, n=3. (E) SPCA2 KD in MCF-10A showed a significant decrease in the cell proliferation compared to control, n=3. (F) Knockdown of SPCA2 in ZR-75 cells was confirmed by qPCR, n=3. (G) SPCA2 KD in ZR-75 showed decrease in cell proliferation compared to control, n=3. (H) SPCA2 overexpression in HS578T increased cell proliferation compared to vector control, n=3. (I) Overexpression of SPCA1 in MCF-7 cells was confirmed by qPCR, n=3. (J) SPCA1 overexpression in MCF-7 did not confer significant decrease in cell proliferation compared to vector control, n=3. (K) Western blot showing SPCA2 D379N overexpression in MDA-MB-231 using Anti-FLAG antibody. (L) Live cell Ca^2+^ imaging traces of Fura2-AM treated MDA-MB-231 cells with or without SPCA2R and SPCA2 D379N (Control n=51 cells, SPCA2R n=50 cells, SPCA2 D379N n=39). Baseline readings in Ca^2+^-free conditions were established for 80 s and then cells were recorded under 2mM Ca^2+^ conditions for 6 minutes and 20 s. Significance: ^ns^ *P* > 0.05, \**P* < 0.05, ** *P* < 0.01, *** *P* < 0.001, **** *P* < 0.0001.

### Loss of SPCA2 activates the ATM/ATR-p53 pathway

As a starting point to understand the signaling mechanisms linking SPCA2 to cell growth and survival, we analyzed the effect of SPCA2 KD on the transcriptome of MCF-7 cells. Consistent with our observations in Fig. 1, top molecular and cellular function categories altered, relative to control, were Cell Death and Survival (*P* value range of 2.44E-06 – 1.00E-22), and Cell Growth and Proliferation (*P* value range of 2.53E-06 – 3.80E-15), with 467 and 495 differentially expressed genes identified, respectively. Among the top canonical pathways identified by Ingenuity analysis, p53 signaling was significantly up regulated (*P* = 3.87E-04) in SPCA2 KD cells (**Fig. 2A-B**). We confirmed mRNA alterations in a representative set of p53 pathway genes by qPCR analysis (**Table S1**).

**Figure 2:**
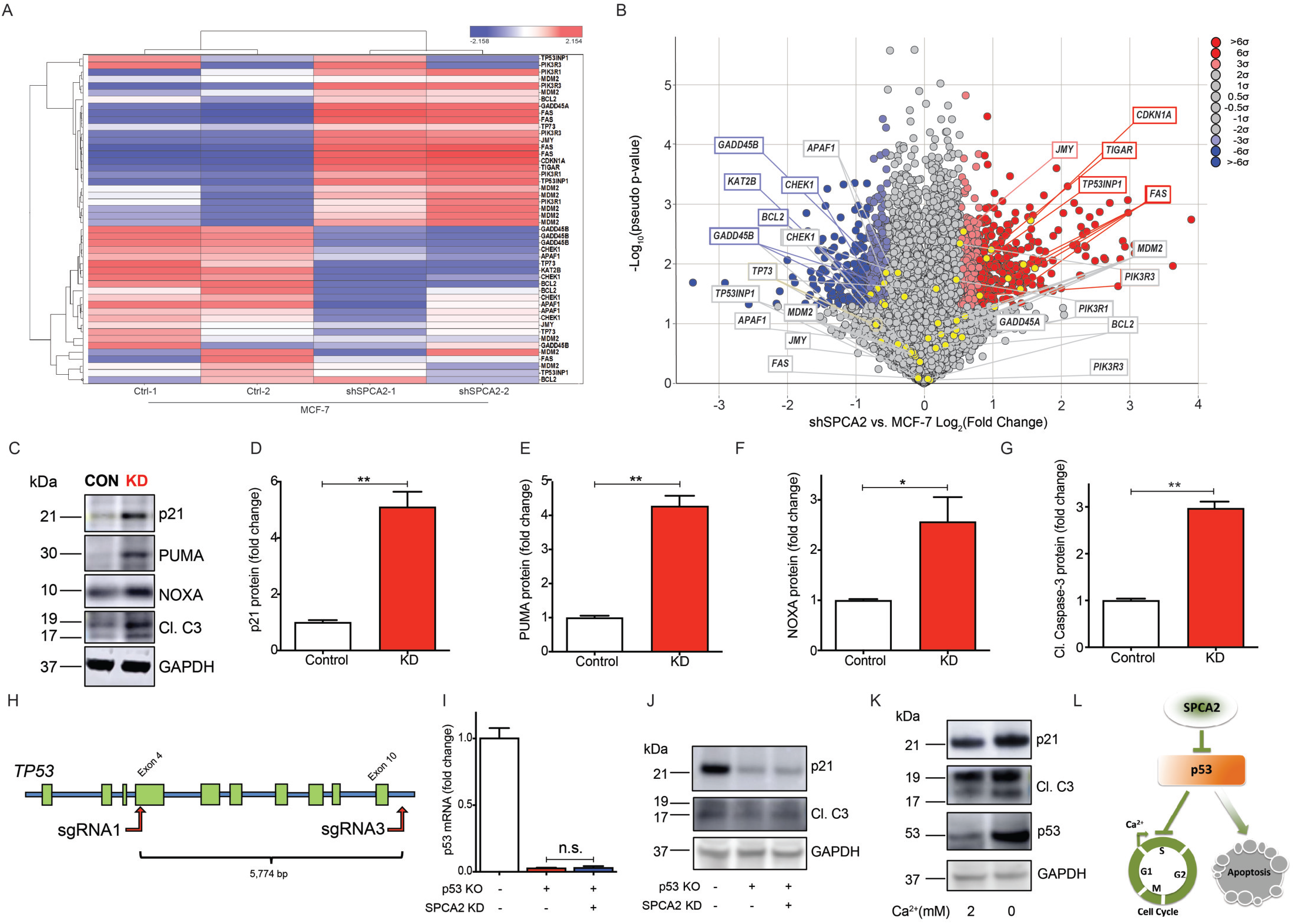
Loss of SPCA2 activates the ATM/ATR-p53 pathway. (A) Heat map with dendrogram indicating differential expression of p53 pathway genes identified from Ingenuity Pathway Analysis. Multiples relate to individual probe sets from the Affymetrix Human Genome U133 Plus 2.0 Array. (B) Volcano plot calculated using pseudo *P* values calculated by oneway ANOVA versus the Log_2_ (fold change). Several p53 pathway genes, indicated in boxes, are highly up (red box) or down regulated (blue box). Color shading correlates with significance of change. (C) Representative western blotting images of p21, PUMA, NOXA, and cleaved caspase-3 in MCF-7 control and SPCA2 KD cells, with GAPDH used as a loading control. (D-G) SPCA2 KD in MCF-7 showed a significant increase in protein level of p53 downstream effector molecules p21, PUMA, NOXA, and cleaved caspase-3 compared to control, n=3. (H) Schematic showing strategy for generating *TP53* gene KO in MCF-7 cells. (I) p53 transcript was not detected in the *TP53* KO MCF-7 cells. (J) SPCA2 KD did not increase p21 and cleaved caspase-3 in *TP53* KO MCF-7 cell line. (K) Brief exposure (2 h) of MCF-7 cells to Ca^2+^-free medium increased expression of p53 and its downstream effectors p21 and cleaved caspase, relative to standard (2 mM Ca^2+^) medium. (L) Schematic showing SPCA2 promotes cell cycle proliferation and blocks apoptosis via p53 pathway. Significance: ^ns^ *P* > 0.05, \**P* < 0.05, ** *P* < 0.01, *** *P* < 0.001.

**Table S1:** Representative set of p53 pathway genes in MCF-7 cells altered by SPCA2KD Comparison of a subset of p53 pathway genes determined to be differentially expressed in the microarray (n = 2) to expression measured by qPCR (n = 3). For genes that contain multiple probe sets in the Affymetrix Human Genome U133 Plus 2.0 Array, a representative value was selected, typically the value of highest significance.

| Gene name | Microarray |  |  |  | qPCR |  |
| --- | --- | --- | --- | --- | --- | --- |
|  | Gene symbol | Probeset ID | Fold Change | Pseudo P-value | Fold Change | P-value |
| Cyclin dependent kinase inhibitor 1A | *CDKN1A* | 202284_s_at | 2.94 | ** | $3.02 \pm 0.48$ | ** |
| Cyclin dependent kinase 4 | *CDK4* | 202246_s_at | -1.14 | * | $-2.09 \pm 0.15$ | * |
| BCL2 associated X, apoptosis regulator | *BAX* | 211833_s_at | 1.29 | * | $1.75 \pm 0.217$ | * |
| BCL2 like 11 | *BCL2L11* | 1558143_a_at | 1.17 | * | $2.32 \pm 0.166$ | * |
| BCL2 binding component 3 | *BBC3* | 211692_s_at | -1.04 | ns | $3.98 \pm 1.02$ | * |
| Phorbol-12-myristate-13-acetate-induced protein 1 | *PMAIP1* | 204285_s_at | 1.36 | ns | $5.01 \pm 1.90$ | * |
| Fas ligand | *FASLG* | 211333_s_at | 1.04 | ns | $5.04 \pm 1.11$ | ** |

\* $p < 0.05$ ; \*\* $p < 0.01$

Activation of p53 plays a critical role in the G0/G1 cell cycle checkpoint arrest by inducing expression of the cyclin-dependent kinase inhibitor p21, and in apoptosis by activation of the pro-apoptotic Bcl-2 family members PUMA and NOXA^22^. Western blot analysis showed that protein levels of p21, PUMA, and NOXA in MCF-7 SPCA2 KD were significantly increased compared to control (**Fig. 2C-F**). Pro-apoptotic proteins activate serine protease caspases, leading to apoptosis^23^. There was a significant increase in cleavage of caspase-3 in SPCA2 KD compared to control (**Fig. 2C and 2G**). Similar results were observed following SPCA2 KD in the ZR-75 cell line (**Figure S2A-B**). In contrast, KD of SPCA1 in MCF-7 did not increase cell cycle or apoptosis markers (**Fig. S2C-D**). To confirm the critical role of p53 in mediating SPCA2 KD phenotypes, we engineered a stable knockout of the *TP53* gene in MCF-7 (**Fig. 2H-I**) and generated SPCA2 KD in this cell line (**Fig. S2E**). In the absence of p53, SPCA2 KD failed to elevate expression of p21, PUMA, NOXA, and cleaved caspase-3 (**Fig. 2J**, **Fig. S2 F-H**).

Previously, we showed that Orai1 was a downstream effector of SPCA2 in mediating SICE in breast cancer cells^14^. Here we show that Orai1 KD in MCF-7 phenocopies SPCA2 KD by similar increases in p21 and NOXA mRNA expression (**Fig. S2I-K**), consistent with previous reports of a role for Orai1 in cell proliferation and apoptosis in cancer cells ^24 25^ Importantly, brief (2 h) removal of external Ca^2+^ from the culture medium, by chelation with EGTA, was sufficient to increase p53 levels, and concomitantly increase p21 and cleaved caspase-3 (**Fig. 2K**). We conclude that store-independent Ca^2+^ entry, activated by SPCA2, exerts a proliferative and pro-survival effect by inhibiting p53 (**Fig. 2L**).

Induction of p53 is mediated mainly through uncoupling p53 from its key negative regulator MDM2, following phosphorylation by the serine/threonine protein kinases ATM/ATR in response to DNA damage^26^. Therefore, we hypothesized a potential link between SPCA2 and DNA damage. There was a significant increase (*P* < 0.001) in nuclear staining and expression of the double-stranded DNA damage marker p-H2AX (**Fig. 3A-C**), as well as the singlestranded DNA damage marker F7-26 (*P* < 0.001) (**Fig. 3D-E**) in SPCA2 KD cells compared to control. Downstream effectors in the DNA damage response pathway were activated in SPCA2 KD cells, as evidenced by the increase (*P* < 0.05) in phosphorylation of ATM, ATR, and p53 (**Fig. 3F-I**). Similar results were observed upon SPCA2 KD in MCF-10A (**Fig. S3A-B**). Following rescue of SPCA2 KD in MCF-7 with a silencing resistant SPCA2R construct or the pump inactive SPCA2 D379N mutant ^10^, we observed no increase of DNA damage, cell cycle arrest, and apoptosis markers (**Fig. S3D-E**).

**Figure 3:**
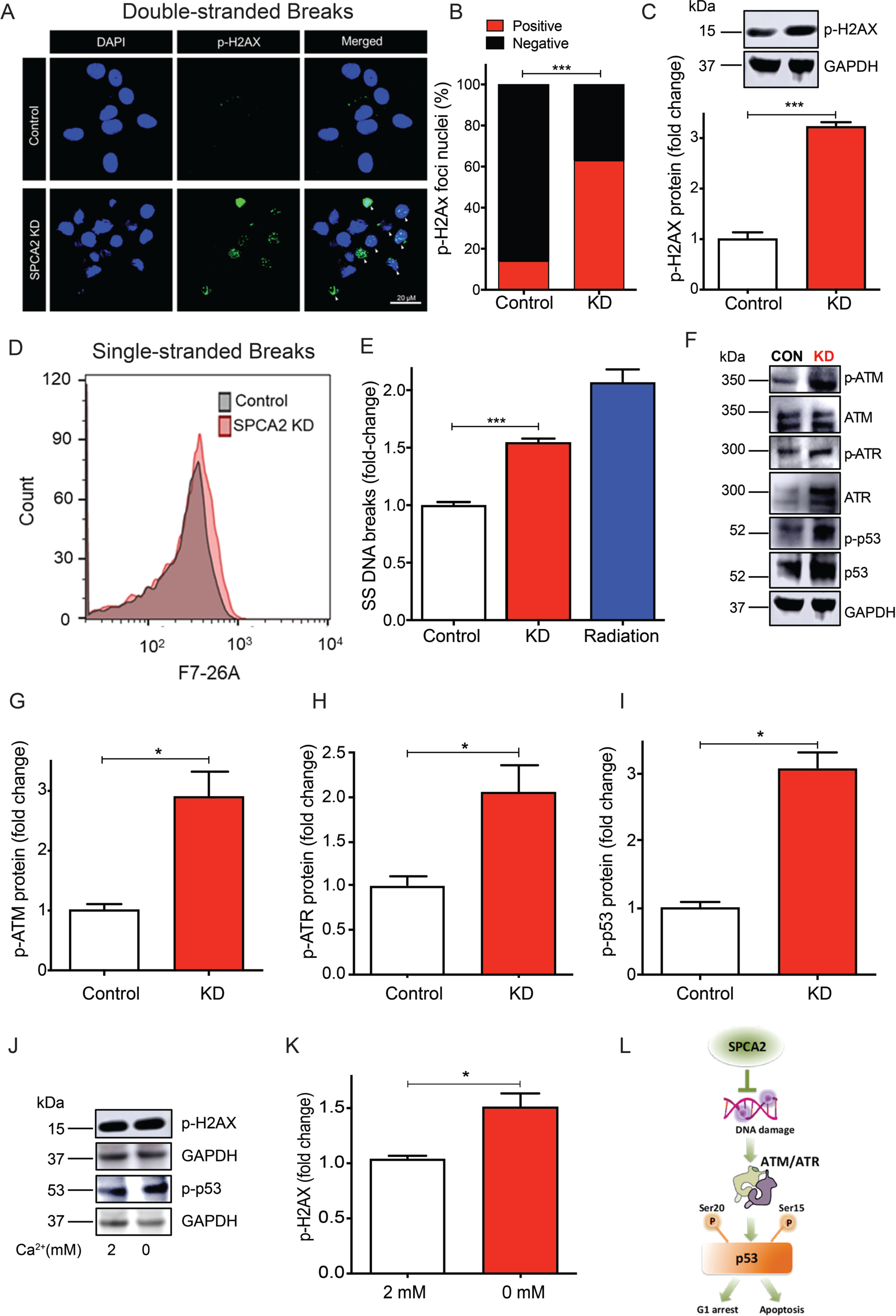
Loss of SPCA2 activates the ATM/ATR-p53 pathway. (A) Representative confocal microscope images showing immunofluorescence staining of p-H2AX (marker for double stranded DNA breaks) in MCF-7 control and SPCA2 KD cells (40x magnification; scale bar, 20 μm). (B) p-H2AX positive and negative nuclei are quantified by ImageJ software. SPCA2 KD showed significant number of p-H2AX positive nuclei compared to control. n=247 in control, n= 250 for SPCA2 KD. (C) p-H2AX protein expression was detected using western blotting; GAPDH was used as a loading control. SPCA2 KD significantly increased p-H2AX expression compared to control, n=3. (D) Representative cell count from flow cytometry showing F7-26 staining (marker for single stranded DNA breaks DNA breaks) in MCF-7 control and SPCA2 KD cells. (E) Single stranded DNA breaks are significantly higher in SPCA2 KD compared to control. Radiation treatment (10 Gy) was used as positive control, n=3. (F) Representative western blotting images of ATM, p-ATM, ATR, p-ATR, p53, and p-p53 in MCF-7 control and SPCA2 KD cells; GAPDH was used as a loading control. (G-I) Densitometry of western blots shows SPCA2 KD significantly increased p-ATM, p-ATR, p-p53 compared to control, n=3. (J) MCF-7 cells were exposed to calcium free media (0 mM) for 1 hour, and cell lysates examined by western blotting. p-H2AX and p-p53 increased relative to cells grown in media with 2 mM calcium. (K) Densitometric analysis shows increased p-H2AX in cells exposed to Ca^2+^ free medium, n=3. (L) Schematic showing SPCA2 blocks DNA damage and downstream activation of ATM/ATR-p53 pathway leading to cell cycle arrest and apoptosis. Significance: \**P* < 0.05, *** *P* < 0.001.

We hypothesized that DNA damage response elicited by loss of SPCA2 was linked to extracellular Ca^2+^. We show that a brief (1 h) exposure of MCF-7 cells to Ca^2+^-free medium significantly increased phosphorylation of H2AX and p53 (**Fig. 3J-K**). These findings reveal an unexpected and novel role for SPCA2-mediated store independent Ca^2+^ entry in maintaining genomic integrity and regulating the DNA damage response (**Fig. 3L**).

**Figure S2:**
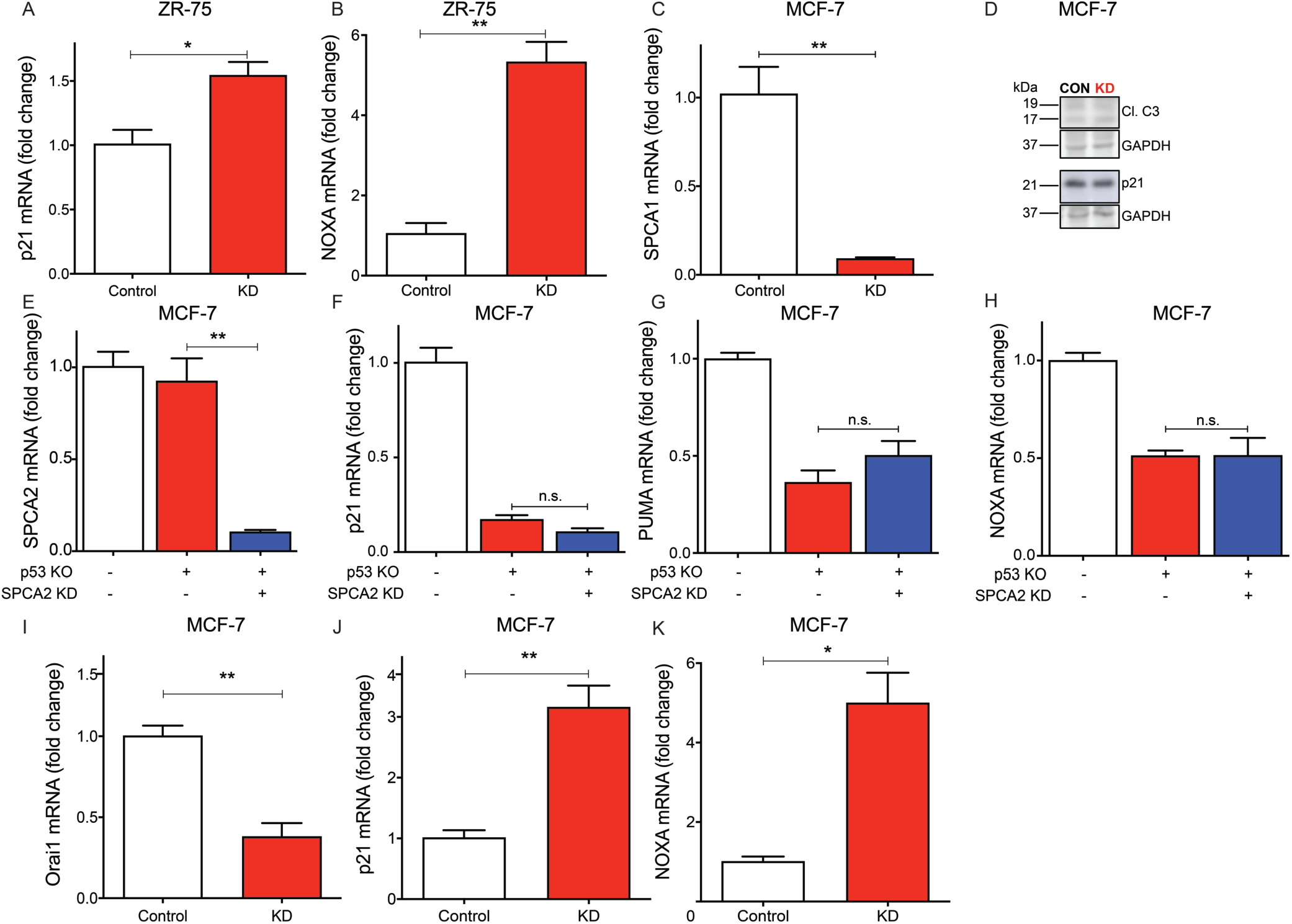
Loss of SPCA2 activates the ATM/ATR-p53 pathway. (A-B) SPCA2 KD in ZR-75 cells significantly increased p21 and NOXA, n=3. (C) Knockdown of SPCA1 was in MCF-7 cells was confirmed by qPCR, n=3. (D) Representative western blotting images of cleaved caspase-3 and p21 in MCF-7 control and SPCA1 KD cells; GAPDH was used as a loading control. (E) SPCA2 KD in p53 KO MCF-7 cell line was confirmed by qPCR, n=3. (F-H) SPCA2 KD did not increase p21, PUMA, and NOXA in p53 KO MCF-7 cell line, n=3. (I) Orai1 knockdown in MCF-7 was confirmed by qPCR, n=3. (J-K) Orai1 knockdown in MCF-7 cells increased p21 and NOXA, n=3. Significance: ^ns^ *P* > 0.05, \**P* < 0.05, ** *P* < 0.01, *** *P* < 0.001.

**Figure S3:**
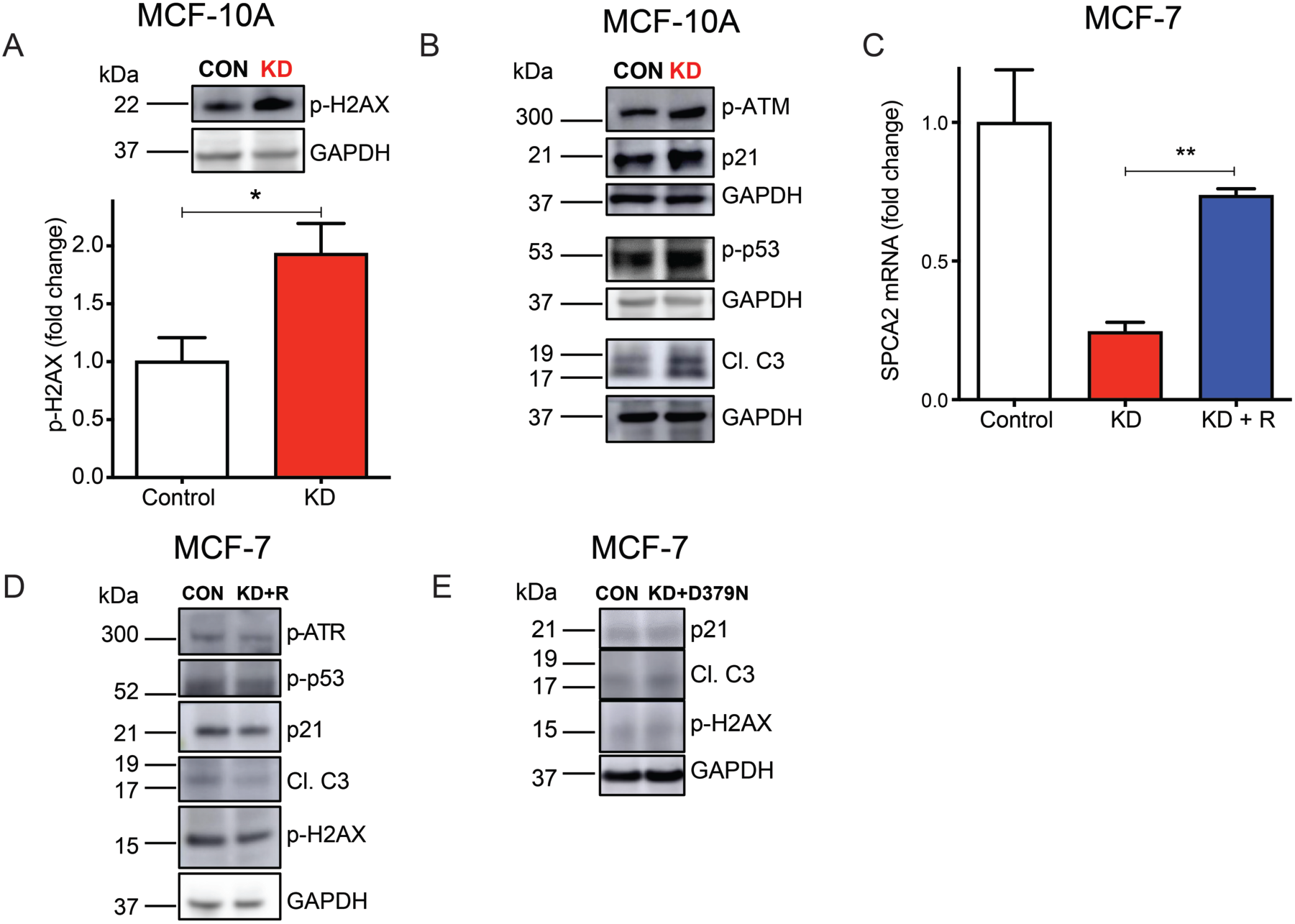
Loss of SPCA2 activates the ATM/ATR-p53 pathway. (A) Representative western blotting images of p-H2AX in MCF-10A control and SPCA2 KD cells; GAPDH was used as a loading control. SPCA2 KD in MCF-10A showed a significant increase in protein level of p-H2AX compared to control, n=3. (B) Representative western blotting images of p-ATM, p21, p-p53, and cleaved caspase-3 in MCF-10A control and SPCA2 KD cells; GAPDH was used as a loading control. (C) SPCA2 transcript was significantly rescued in MCF-7 cells after knockdown (KD), by recombinant, silencing-resistant SPCA2R construct (R) as confirmed by qPCR, n=3. (D) Representative western blotting images of p-ATM, p-p53, p21, cleaved caspase-3, p-H2AX in MCF-7 control and SPCA2 KD rescued cells (KD+R); GAPDH was used as a loading control. (E) Representative western blotting images of p21, cleaved caspase-3, p-H2AX in MCF-7 control and SPCA2 KD rescued with silencing resistant, recombinant SPCA2R carrying the loss of function D379N mutation (KD+D379N); GAPDH was used as a loading control. Significance: \**P* < 0.05, ** *P* < 0.01.

### SPCA2 protects against ROS-mediated DNA damage

Intracellular reactive oxygen species (ROS) are known to initiate DNA damage and activate the ATM/ATR-p53 pathway^27^. Therefore, we evaluated the effect of SPCA2 on ROS production. SPCA2 KD significantly increased (*P* < 0.001) endogenous levels of ROS, as measured by DCFDA fluorescence, in MCF-7 (**Fig. 4A-B**) and MCF-10A (**Fig. S4A-B**). In response to oxidative stress induced by treatment with H2O2, ROS levels were further elevated, with SPCA2 KD cells showing greater increase (*P* < 0.05) compared to control (**Fig. 4A-B**, **S4A-B**). Similarly, brief removal (10 min) of extracellular Ca^2+^ significantly increased ROS compared to cells maintained in 2 mM calcium, and the absence of Ca^2+^ exacerbated ROS production in the presence of H2O2 (**Fig. 4C-D**). Mitochondria are one the principle generators of ROS, through activity of the electron transport chain^28^. We observed a significant increase (*P* < 0.05) of mitochondrial ROS (MitoSOX-Red) in SPCA2 KD compared to control (**Fig. 4E-F**). Doxorubicin was previously reported to induce mitochondrial ROS^29^. Upon doxorubicin treatment, SPCA2 KD showed greater increase of (*P* < 0.01) of mitochondrial ROS compared to control (**Fig. 4E-F**). As expected, following rescue of the MCF-7 SPCA2 KD with silencing resistant SPCA2R construct, we observed no increase of ROS (**Fig. S4C-D**) (*P* > 0.05).

**Figure 4:**
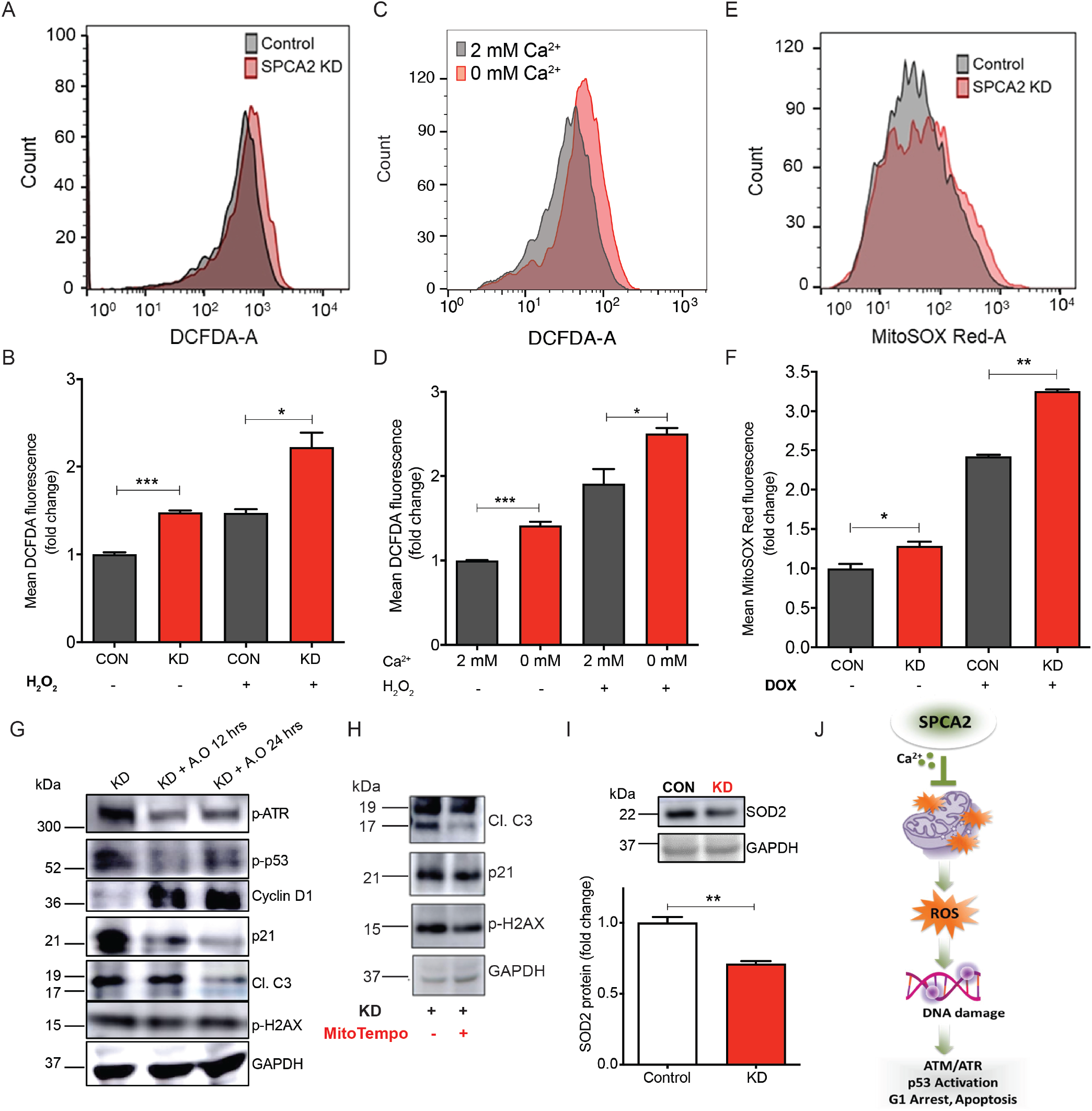
SPCA2 protects against ROS-mediated DNA damage. (A) Cell count distribution from representative flow cytometry analysis measuring ROS (DCFDA fluorescence) in MCF-7 control and SPCA2 KD cells. (B) ROS production was significantly increased compared to control. H_2_O_2_ (500 μM for 20 minutes) was used as positive control; n=6 for SPCA2 KD and control; n=3 for H_2_O_2_ experiment. (C) Representative flow cytometry showing ROS (DCFDA) in MCF-7 cells exposed to Ca^2+^-free media for 10 min compared to cells grown in media with 2 mM Ca^2+^. (D) ROS production increased in Ca^2+^-free media; H_2_O_2_ (500 μM for 15 minutes) was used as positive control, n=3. (E) Representative flow cytometry showing mitochondrial ROS (MitoSOX-Red) of MCF-7 control and SPCA2 KD cells. (F) Mitochondrial ROS was significantly increased in SPCA2 KD compared to control. Doxorubicin was used as positive control (500 nM for 24 hours), n=3. (G) Antioxidants (Vitamin C, 500 μM + N-acetyl cysteine, 500 μM) were added to MCF-7 SPCA2 KD cells for 12 hours and 24 hours and immunoblotting is performed using GAPDH as loading control. Anti-oxidants decreased p-ATM, p-p53, p21, cleaved caspase-3, p-H2AX, and increased cyclin D1. (H) Mitochondrial anti-oxidant (MitoTempo, 5 μM) was added to MCF-7 SPCA2 KD cells for 24 hours and immunoblotting was performed using GAPDH as loading control. MitoTempo decreased levels of cleaved caspase-3, p21, and p-H2AX. (I) SOD2 protein expression was detected using western blotting, and GAPDH was used as a loading control. SPCA2 KD significantly decreased SOD2 expression compared to control, n=3. (J) Schematic showing SPCA2 blocks ROS mediated activation of DNA damage, activation of ATM/ATR-p53 pathway, resulting in cell cycle arrest and apoptosis. Significance: \**P* < 0.05, ** *P* < 0.01, *** *P* < 0.001.

Addition of anti-oxidants (N-acetyl cysteine + vitamin C) known to block ROS, decreased the activation of DNA damage response in the ATM/ATR-p53 mediated pathway, as seen by decrease in phosphorylated ATR, p53 and p21, increased expression of Cyclin D1, decreased production of cleaved caspase-3 and decreased levels of p-H2AX in SPCA2 KD cells (**Fig. 4G**). Similarly, addition of mitochondrial-specific antioxidant Mito-TEMPO to SPCA2 KD reduced cleaved caspase 3, p21 expression and p-H2AX levels in MCF-7 KD cells (**Fig. 4H**). We observed a significant decrease in mitochondrial-specific manganese superoxide dismutase (MnSOD/SOD2) expression in the SPCA2 KD cells compared to control (**Fig. 4I**), suggesting a disruption of ROS scavenging mechanisms in mitochondria. Taken together, these results reveal a novel role for SPCA2 in protection against mitochondrial ROS mediated DNA damage and activation of ATM/ATR-p53 pathways (**Fig. 4J**).

**Figure S4:**
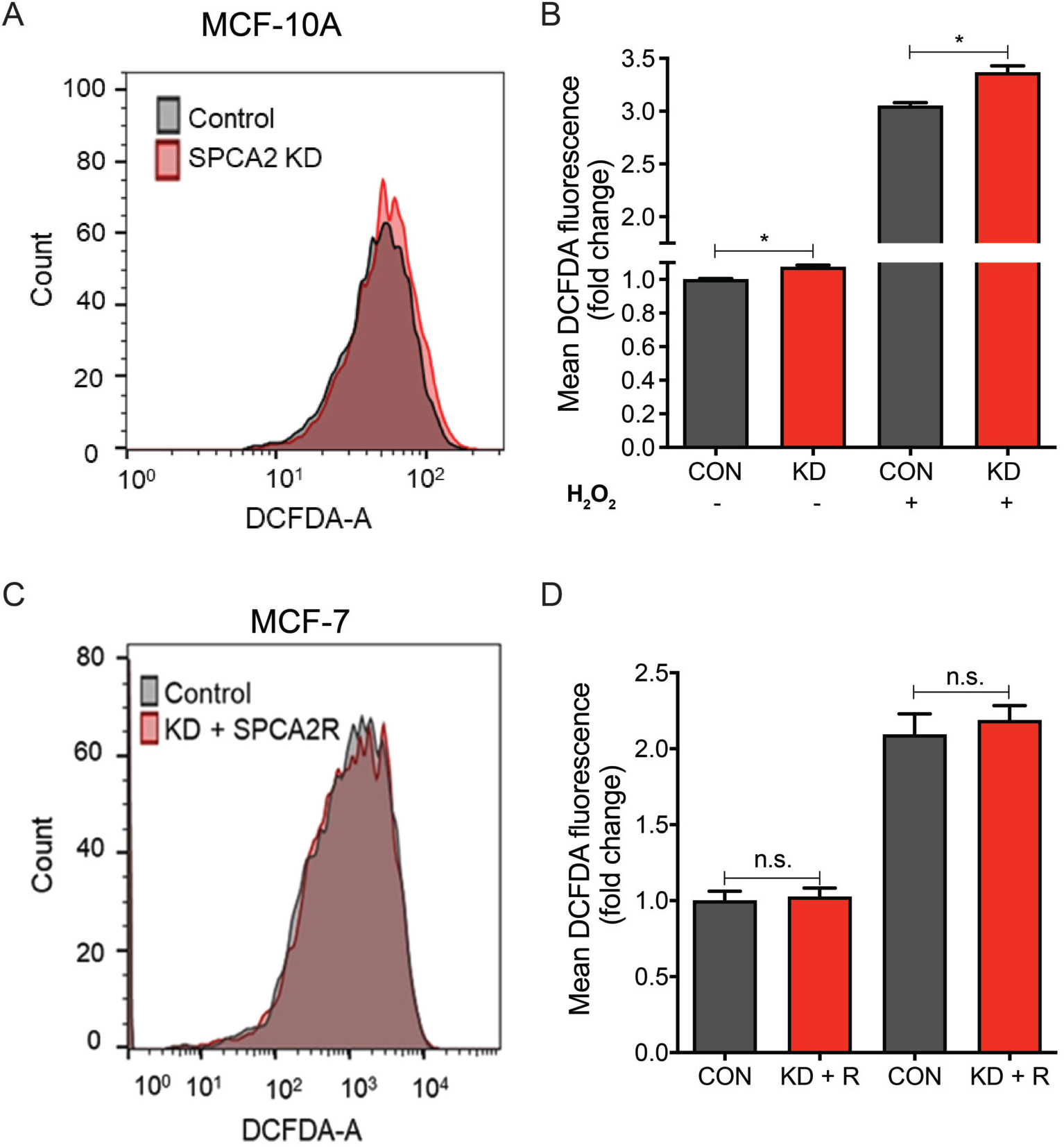
SPCA2 protects against ROS-mediated DNA damage. (A) Representative flow cytometry showing ROS production in MCF-10A control and SPCA2 KD cells. (B) ROS production is significantly increased in SPCA2 KD compared to control. H_2_O_2_ (500 μM for 20 minutes) was used as positive control. n=3 for SPCA2 KD and control, n=2 for H_2_O_2_ experiment. (C) Representative flow cytometry showing ROS production in MCF-7 control and SPCA2 KD rescued (KD + SPCA2R) cells. (D) There was no significant difference in ROS production between control and SPCA2 KD rescued cells. n=3. Significance: ^ns^ *P* > 0.05, \**P* < 0.05

### Loss of SPCA2 expression sensitizes breast cancer cells to DNA damaging agents

Drugs that elicit the DNA damage response, including carboplatin, doxorubicin and ionizing radiation, are used as neoadjuvant therapy for breast cancer patients^30, 31^. Since SPCA2 KD increased DNA damage and activation of ATM/ATR-p53 pathway, we investigated if loss of SPCA2 expression sensitizes breast cancer cells to DNA damaging agents. We show that MCF-7 SPCA2 KD cells are more sensitive to carboplatin, doxorubicin and ionizing radiation, compared to control (**Fig. 5A-C**). Similarly, SPCA2 KD conferred doxorubicin sensitivity to noncancer MCF-10A cells (**Fig. S5A**). In contrast, there was no significant difference in response to non-DNA damaging chemotherapeutic agents used to treat breast cancer such as paclitaxel (anti-microtubule agent), methotrexate (anti-folate) and tamoxifen (estrogen blockage) in SPCA2 KD cells as compared to control (**Fig. 5D-F**). As expected, rescue of knockdown using the silencing resistant SPCA2R construct restored doxorubicin resistance to MCF7 SPCA2 KD cells (**Fig. S5B**).

**Figure 5:**
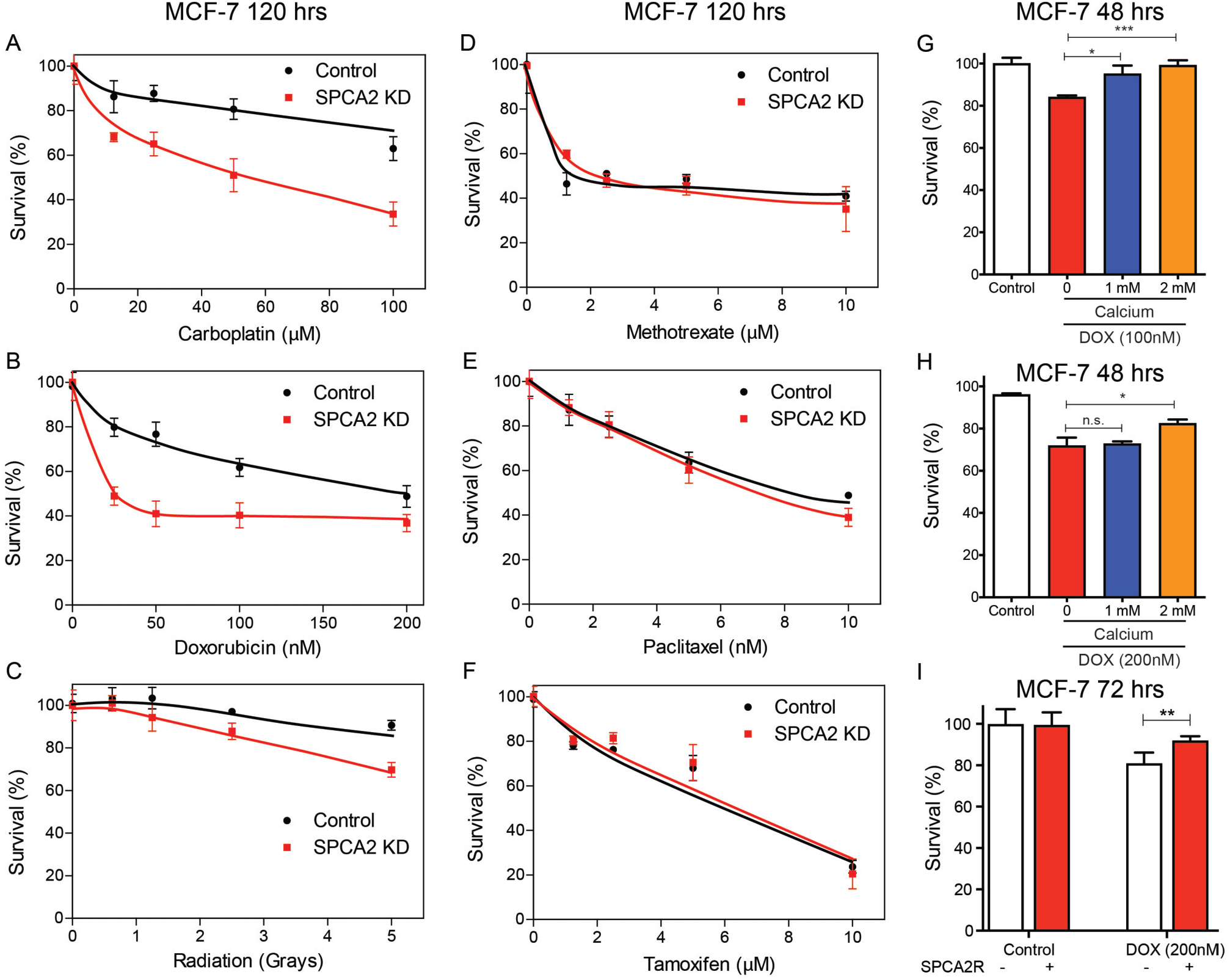
Loss of SPCA2 expression sensitizes breast cancer cells to DNA damaging agents. Dose-response curves of control and SPCA2 KD in MCF-7 cells. Error bars represent standard deviation (n = 5 or 6). Approximately 3,000 – 5,000 cells were seeded into 96 well plates, and incubated overnight. Cells were treated with drugs 120 hours before MTT cytotoxicity analysis. The survival fraction was determined by mean fluorescence of the treated cells/mean fluorescence of control cells, and normalized to untreated. (A-C) SPCA2 KD increased chemosensitivity to the DNA damaging agents doxorubicin (0-200 nM), carboplatin (0-100 μM), and radiation (0-5 grays). (D-F) SPCA2 KD did not increase chemosensitivity to the non-DNA damaging agents to methotrexate (0-10 μM), paclitaxel (0-10 nM), and tamoxifen (0-10 μM). (G-H) Approximately 10,000 cells were seeded into 96 well plates in Ca^2+^-free media supplemented with 2% FBS without addition of EGTA, and incubated overnight. Cells were treated with doxorubicin (100 and 200 nM) for 48 hours, along with/without addition of Ca^2+^ (1 mM and 2 mM). Cells cultured in Ca^2+^-free media were considered as control. Culturing MCF-7 cells with increasing concentration of Ca^2+^ significantly rescued doxorubicin cytotoxicity. n=5 for 100 nM, n=4 for 200 nM. (I) Approximately 10,000 cells were plated and incubated overnight. Cells were then treated with doxorubicin (DOX; 200 nM). Chemosensitivity to doxorubicin is significantly reversed by forced overexpression of SPCA2R, n=5. Significance: ^ns^*P* > 0.05, \**P* < 0.05, \*\**P* < 0.01.

Since removal of extracellular Ca^2+^ or knockdown of SPCA2 increased ROS mediated DNA damage, leading to increased sensitivity to DNA damaging agents, we asked if doxorubicin cytotoxicity was modulated by extracellular calcium. We show that culturing MCF-7 cells in increasing concentrations of calcium significantly rescued doxorubicin cytotoxicity (*P* < 0.05) (**Fig. 5G-H**). Furthermore, ectopic expression of SPCA2 in MCF-7 (SPCA2R; **Fig. S5C**) was found to significantly blunt doxorubicin cytotoxicity (**Fig. 5I**). These results point to SPCA2 as a novel chemoresistance mechanism that links Ca^2+^ homeostasis to the response to DNA damaging chemotherapeutic drugs.

**Figure S5:**
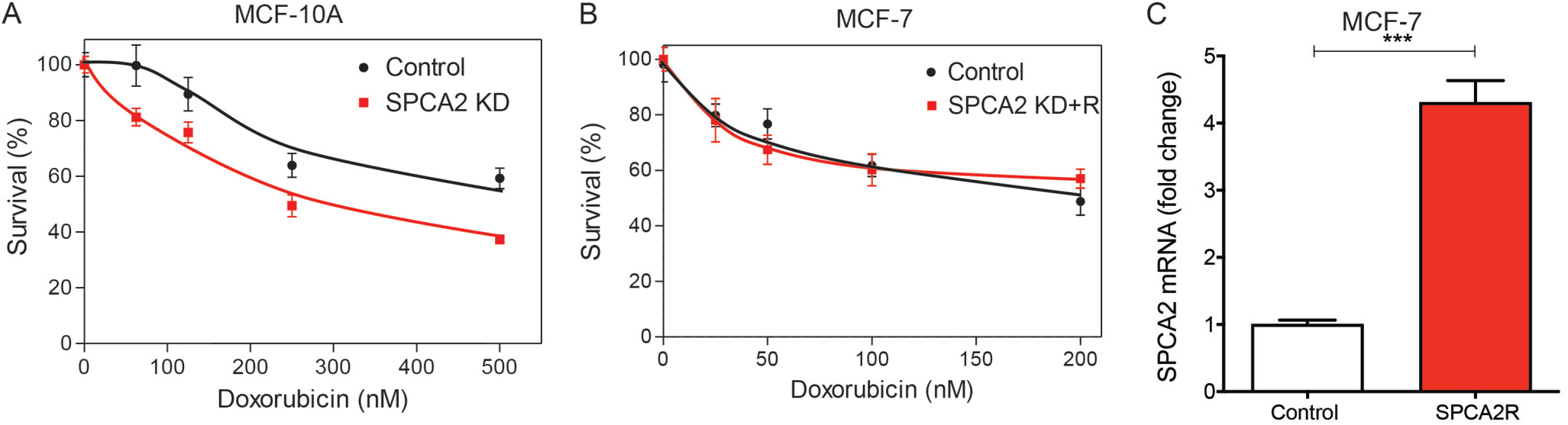
Loss of SPCA2 expression sensitizes breast cancer cells to DNA damaging agents. (A) SPCA2 KD in MCF-10A cells increased chemosensitivity to doxorubicin (0-500 nM) compared to control (n=5 or 6, time=120 hours). (B) SPCA2 KD rescued cells in MCF-7 cells did not increase chemosensitivity to doxorubicin (0-200 nM) compared to control (n=5 or 6, time=120 hours). (C) Lentiviral mediated expression of recombinant SPCA2 (SPCA2R) in MCF-7 cells resulted in significant increase in transcript levels, n=3. Significance: *** *P* < 0.001.

## DISCUSSION

### Multiple mechanisms of Ca^2+^ entry drive cell cycle progression in cancer cells

Aberrant Ca^2+^ signaling and homeostasis impacts all aspects of breast cancer, ranging from cell proliferation, survival, and metabolism to chemoresistance and death. Key regulators of Ca^2+^ homeostasis, including ion channels and pumps, show subtype-specific dysregulation in breast cancer, offering the potential of tailored intervention as a therapeutic option^2, 9, 32^. The secretory pathway Ca^2+^, Mn^2+^-ATPase Isoform 2 (SPCA2) is prominently elevated in luminal breast cancer subtypes that express ER, PR or HER2 receptors where it interacts with plasma membrane ion channels to elicit constitutive, store-independent Ca^2+^ entry ^2, 18^. Ca^2+^ influx is critical for several steps in cell cycle progression, including the initiation of DNA synthesis and entry into S-phase^33^. The absence of STIM1 and Orai1 or Orai3, mediators of store operated Ca^2+^ entry (SOCE), blocked G1/S transition in cancer cells^20, 34 21^. Here, we show that store independent Ca^2+^ entry (SICE) mediated by SPCA2 is important for cell cycle progression and survival by promoting progression through the G1/S checkpoint in breast cancer cells. Thus, multiple mechanisms of Ca^2+^ entry are involved in driving cancer cell proliferation, depending on the subtype specific expression of Ca^2+^ regulators and channels.

### Emerging links between ion homeostasis and p53 signaling

As evidenced by our transcriptomic analysis, cell cycle arrest and apoptosis in the absence of SPCA2 was mediated by the tumor suppressor p53 pathway, which activates well-known downstream effectors including p21, NOXA and PUMA. Similarly, silencing the Orai1 Ca^2+^ channel also elicits p21 expression (**Fig. S2H-J**), and conversely, overexpression of Orai3 decreases transcriptional activity of p53 in breast cancer cells^35^. These findings reveal a link between Ca^2+^ entry and p53 mediated cell cycle arrest and apoptosis. The specific molecular pathways connecting Ca^2+^ entry to p53 function are complex and not completely understood. For example, a Ca^2+^-dependent and transcription-independent role of p53 in regulating apoptosis at the endoplasmic reticulum has been reported^36^. Hasna et al. have proposed that Ca^2+^ entry through Orai3 in breast cancer cells signals through the PI3K/Sgk-1/Sek-1 kinases to activate the ubiquitin ligase Nedd4-2, targeting p53 for degradation^35^. Here, we demonstrate that p53 regulation is (i) independent of pumping activity of SPCA2, (ii) linked to SICE in breast cancer cells and (iii) occurs by upstream regulation of the DNA damage response pathway.

This study also reveals an unexpected role for SPCA2 in mitochondrial integrity and function that warrants further study. We observed decreased expression of the mitochondrial-specific manganese superoxide dismutase (MnSOD/SOD2), which could account for increased mitochondrial ROS accumulation in the absence of SPCA2. We note that tumor cells also accumulate Mn^2+^, which is a Ca^2+^ surrogate that can gain entry through SICE^14, 37^. Thus, dysregulation of both cellular Ca^2+^ and Mn^2+^ homeostasis by SPCA2 could alter p53 function in breast cancer cells.

### Importance of the SPCA2 – p53 axis on chemotherapy response in breast cancer

More than two-third of all breast cancers are ER+, and SPCA2 is up regulated in a majority of these tumors^10^. Although p53 is mutated in >50% of all cancers, where it contributes to transformation, the majority (70-80%) of breast cancers are p53^WT^. Specifically, only 17% of luminal A (ER+, PR+/-, HER2-) and 41% of luminal B (ER+, PR+/-, HER2+/-) carry p53 mutations, whereas p53 mutations occur mostly in basal, ER-tumor types (e.g., MDA-MB-231 cell line)^38^. The mutation status of p53 is especially relevant for the clinical treatment of ER+ breast cancer, which often involves chemotherapy followed by adjuvant therapy consisting of tamoxifen or aromatase inhibitors.

Chemotherapy includes DNA damaging drugs like doxorubicin, which induce p53 mediated apoptotic response. Paradoxically, ER+ breast cancer cells are relatively insensitive to chemotherapy-induced apoptosis, even in the presence of p53^WT^. Bailey et al. showed that ER blocked the induction of a subset of pro-apoptotic genes by p53, resulting in repression of the apoptotic response^39^. Consistent with these findings, ER+ p53^WT^ tumor cells show hallmarks of senescence and dormancy, followed by further tumor proliferation after the end of treatment^38^. In contrast, genetic abnormalities accumulate in p53-mutated tumors, resulting in mitotic catastrophe with further response to treatment. Taken together, it is clear that p53^WT^ tumors have poorer response to alkylating agents in the clinic that needs to be addressed.

In MCF-7 cells, doxorubicin treatment activates p53 to mediate cell death, and there is less p53 activation in doxorubicin resistant cells^40^. We show that Ca^2+^ entry negatively regulates the p53 pathway, either by targeting p53 for degradation or inhibiting upstream activating mechanisms as reported in this study. Thus, SPCA2 knockdown triggers robust p53 activation in ER+ p53^WT^ breast cancer cells resulting in enhanced sensitivity to DNA damaging agents, including doxorubicin. Our findings confirm and mechanistically extend a previous report showing that increasing levels of external Ca^2+^ confer doxorubicin resistance in MCF-7 cells^41^. Furthermore, we showed that mitochondrial ROS generated by doxorubicin is exacerbated in SPCA2 KD cells and accompanied by decreased levels of MnSOD. In this context, high expression of MnSOD was shown to promote survival of circulating breast cancer cells and increase resistance to doxorubicin^42^.

Pharmacological targeting of the DNA damage response pathway is of growing interest in cancer therapy^43, 44^. We show that store independent Ca^2+^ entry is required for genomic stability and contributes to resistance against DNA damaging drugs. Taken together with clinical data showing poor survival prognosis for ER+/PR+ breast cancer patients expressing high SPCA2, our findings should motivate a search for drugs that target components of store-independent Ca^2+^ entry pathway in hormone receptor positive breast cancer cells and potentially synergize with DDR pathway inhibitors.

## METHODS AND MATERIALS

### Cell culture

Cells used in this study were MCF-7 (ATCC HTB-22), MCF-10A (ATCC CRL-10317), MDA-MB-231 (ATCC HTB-26), Hs578T (ATCC HTB-126), and ZR-75 (gifted by Dr. Nakshatri, Indiana University School of Medicine). MCF-7, MDA-MB-231 and Hs578T cells were gown in DMEM (Thermo Fisher Scientific, Waltham, MA) containing antibiotic, and 5-10% FBS (Sigma Aldrich, St. Louis, MO). For calcium free experiments, cells were grown in calcium free DMEM, with 2% FBS and 150 μM EGTA ^35^. MCF-10A was cultured in DMEM/F12 containing 5% horse serum, 20 ng/mL EGF, 0.5 mg/mL hydrocortisone, 100 ng/mL cholera toxin, 10 μg/mL insulin, and 1X penicillin/streptomycin. ZR-75 was cultured RPMI 1640 containing antibiotic, and 10% FBS. Cells were cultured with 5% CO_2_ and at 37°C in a humidified incubator. All cell lines were grown for less than a month or no more than 6 passages. We did not see visible contamination by mycoplasma. Since SPCA2 expression is regulated by cell density^18^, cells were maintained at 50-60% confluency in both knockdown experiments and controls. Although we used lentiviral transfection, resistant clones tended to appear after 2-3 passages, therefore all experiments used early passage cells.

### Lentiviral Transfection

The silencing resistant SPCA2R construct has been previously described^10, 18^ FUGW overexpression constructs and pLK0.1 shRNA and lentiviral construction of both SPCA isoforms were packaged and transfected according to previous methods^18^ using pCMV-Δ8.9 and PMDG at a ratio using 9:8:1 in HEK293T cells. Virus was collected after 24 hours for 3 consecutive days and concentrated with Lenti-X Concentrator (Takara Bio USA, Mountain View, CA). A mixture of two shRNA constructs for SPCA2 was used. Cells were transfected with virus for 48 hours, and selected with puromycin at varying concentrations according to kill curves performed for each specific cell line (0.5-2 μg/mL). Knockdowns were confirmed by qPCR. Experiments were performed within 2-3 passages to ensure that knockdown was maintained.

### *TP53* knockout in MCF-7 cells

TP53 knockout MCF-7 cells were generated as previously published ^45^. Human codon-optimized *Streptococcus pyogenes* wild type Cas9 (Cas9-2A-GFP) was obtained from Addgene (Cambridge, MA). Chimeric guide RNA expression cassettes with different sgRNAs (sgRNA1: CCATTGTTCAATATCGTCCG; sgRNA3: TGGTTATAGGATTCAACCGG) were ordered as gBlocks. These gBlocks were amplified by PCR using primers: gBlock_Amplifying_F: 5’-GTACAAAAAAGCAGGCTTTAAAGG-3’ and gBlock_Amplifying_R: 5’-TAATGCCAACTTTGTACAAGAAAGC-3’. The PCR product was purified by Agencourt Ampure XP PCR Purification beads per the manufacturer’s protocol (Beckman Coulter, Brea, CA). One microgram of Cas9 plasmid and 0.3 μg of each gRNA gBlock (pair: sgRNA1 & sgRNA3) were cotransfected into MCF-7 cells via Lipofectamine 3000 in a 6-well plate. Knockout cells created using the pair sgRNA1 & sgnRNA3 were named KO3.4. The knockout pool was cultured in 10 μM Nutlin-3a (SelleckChem, Houston, TX) for 2 months, changing nutlin-3a treated media every 3 days and passaging cells every 6 to 8 days. The isogenic clone KO3.4E1 was isolated from the knockout pool via limiting dilution in a 96-well plate and incubated at 37 °C in a CO_2_ incubator for 15 days.

DNA was isolated from each MCF-7 wild type and knockout single cell clones following the Agencourt DNAdvance genomic DNA isolation kit. MCF-7 KO3.4E1 cells were PCR amplified using primers: TP53_exon_4_F and TP53_Woke_R: 5’-ATTAGGCCCCTCCTTGAGAC-3’. Products were sent to Eton Bioscience Inc. who purified the PCR products and performed Sanger Sequencing.

### Cell Proliferation

MTS growth assays were performed on adherent cells by plating 1000-5000 cells per well in a 96-well plate and assaying using CellTiter 96^®^ AQueous One Solution (Promega, Madison, WI) and incubating for 2 hours and reading at 490nm. For MCF-10A and ZR-75 knockdown experiment, 0.3 million cells were platted in a 6 well dish and cell proliferation was measured using trypan blue (Invitrogen, Carlsbad, CA) staining.

### Cell Cycle Analysis

Cell cycle analysis was performed by dissociating adherent cells and fixing them in 4% paraformaldehyde for 30 minutes on ice. For better nuclear staining, fixed cells were then frozen for 1 to 2 weeks and thawed on ice and washed with PBS. Cells were treated with 100ug/ml of RNAse A (Thermo Fisher Scientific) and stained with 50ug/ml propidium iodide (Invitrogen) prior to running flow cytometry via FACSAria (BD Biosciences, San Jose, CA).

### cDNA synthesis & Quantitative PCR

1μg of RNA was collected and used for cDNA synthesis (Applied Biosystems, Foster City, CA). The qPCR mastermix was obtained from Thermo Fischer Scientific, 50ng of cDNA, and Taqman probe (Thermo Fischer Scientific) as specified; GAPDH (Hs02796624_g1), SPCA2 (Hs00939492_m1), SPCA1 (Hs00995930_m1), PUMA (Hs00248075_m1), p21 (Hs00355782_m1), and NOXA (Hs00560402_m1).

### Apoptosis assays

#### TUNEL assay

Apoptosis was measured using terminal deoxynucleotidyl transferase dUTP nick end labeling (TUNEL) assay (APO-DIRECT Kit, BD Biosciences). After treatment, cells were fixed in 1% (w/v) paraformaldehyde (Affymetrix, Santa Clara, CA) in PBS and stored in 70% ethanol at −20°C for 24 hours. Cells were rinsed with 1X PBS and incubated with 50 μL of the DNA labeling solution for three hours at 37°C. After incubation, cells were washed twice and suspended in 500 μL of propidium iodide/RNase staining buffer for 30 minutes of counterstaining, then analyzed with flow cytometry (BD LSR-II, BD Biosciences). Data analysis was performed using FlowJo software (Ashland, OR) and plotted using GraphPad Prism (La Jolla, CA).

#### Annexin V

According to the manufacturer’s protocol, cells were trypsinized and washed with PBS. Cells were then washed in annexin binding buffer (ThermoFisher) and stained with annexin V for 15 minutes. Stained cells were then diluted 1:5 in binding buffer and analyzed by flow cytometry using FACSAria.

#### Immunofluorescence of Cleaved caspase-3

Cells were cultured on glass coverslips and were rinsed with PBS and pre-extracted with 1X PHEM buffer, 8% sucrose and 0.025% saponin. Cells were fixed with 4% paraformaldehyde for 30 minutes and were rinsed and washed with PBS 3 times for 5 minutes each. After blocking in 1% BSA, cells were incubated with overnight in 4°C using cleaved caspase-3 (Asp175) Antibody (Cell Signaling Technologies, Danvers, MA). Cells were rinsed with 0.2% BSA 3 times for 5 minutes and were then incubated with a fluorescent secondary antibody in 1% BSA and 0.025% saponin buffer for 30 minutes at room temperature. Coverslips were washed and mounted onto slides with mounting media (Agilent, Santa Clara, CA). Immunofluorescent staining was analyzed using ImageJ.

### Western blotting

Cells were collected, and lysed in RIPA buffer (Thermo Fisher Scientific) supplemented with protease inhibitor cocktail, and phosphatase inhibitors. Protein quantification was done by the bicinchoninic acid assay. Thirty μg of protein in each sample was resolved by electrophoresis using 4-12% bis-tris gels (Thermo Fisher Scientific) and transferred to nitrocellulose membrane (Bio-Rad, Hercules, CA). Immunoblots were probed with antibodies (1:1000) from Cell Signaling Technologies, followed by incubation with HRP-conjugated secondary antibodies. Proteins were visualized using chemiluminescence substrate. Blots were analyzed using ImageJ.

### Live cell calcium imaging

Live imaging of Ca^2+^ was performed using Fura2-AM (Invitrogen) ^18^. MDA-MB-231cells with or without SPCA2R and SPCA2D379 (Control n=51cells, SPCA2R n=50 cells, SPCA2D379=39) were treated with Fura2-AM in imaging buffer (20 mM Hepes, 126 mM, NaCl, 4.5 mM KCl, 2 mM MgCl2, 10 mM glucose at pH7.4) for 30 min. Cells were excited at 340 nm and 380 nm, and Fura emission was captured at 505 nm. To show store independent Ca^2+^ entry (SICE), cells were briefly washed in nominally Ca^2+^-free buffer followed by addition of Ca^2+^ (2mM). Baseline readings in calcium-free conditions were established for 80 sec and then cells were recorded under 2mM Ca^2+^ conditions for 6 minutes.

### Microarray analysis

RNA was isolated using QIAGEN’s RNEasy Mini (Qiagen, Germantown, MD) manufacturer’s protocol. Samples were submitted to the Johns Hopkins Transcriptomics and Deep Sequencing Core for expression analysis. The mRNA concentrations and integrity were determined with Agilent’s NanoDrop instrument then amplified and biotinylated with the Affymetrix GeneChip 3’ IVT PLUS Reagent kit (ThermoFisher Scientific) following manufactirer’s protocol. The labeled samples were then hybridized to Affymetrix Human Genome U133 Plus 2.0 Array GeneChips (ThermoFisher Scientific, Waltham MA) and scanned Using Affymetrix’ GeneChip Scanner 3000 7G and default parameters. Raw data generated as CEL files by the Affymetrix Expression Console were imported for normalization and analysis into Partek Genomics Suite v7.0 (Partek Inc. St Louis MO, USA). Data were extracted using the Affymetrix na36 transcript annotation, log_2_ transformed, and quantile normalized with the RMA (Robust Multi-Array Average) algorithm. This yielded data for 44.6K transcripts, 21K unique NCBI Entrez genes, whose annotation was subsequently updated to current HGNC/NCBI nomenclature. After data quality control the samples’ gene expression were independently compared using a two tailed one way ANOVA between their shSPCA2_1Way vs. control MCF7 biological classes, which provided relative levels of gene expression and statistical significance, as fold change and imputed p-value respectively. Genes whose log_2_ fold changes were greater than 2SD from the mean were deemed to be differentially expressed.

### Detection of DNA damage

#### Single stranded DNA damage

Cells were fixed in methanol-PBS (6:1) and analyzed 1–3 days after fixation. Staining with F7-26 monoclonal antibody (Millipore Sigma, Burlington, MA) was performed according to manufacturer’s instructions^46^: Fixed cells were resuspended in 0.25 ml of formamide and heated in a water bath at 75°C for 10 min. The cells were then returned to room temperature and washed with 2 ml of 1% nonfat dry milk in PBS for 15 min, resuspended in 100 μl of anti-ssDNA Mab F7-26 (100 μg (0.5 ml) antibody in 4.5 ml of 5% FBS in PBS) or 100 μl of isotype control (IgM) Mab (10 μg/ml), incubated at room temperature for 25 min, washed with PBS, and then stained with 100 μl of fluorescence-conjugated goat antimouse IgM antibody (1:50 in 1% nonfat dry milk in PBS) for 25 min at room temperature. Cells were filtered 0.7 micrometer cell filter to filter the cell debris. Cells were then washed with PBS and resuspended in 0.5 ml of PBS containing 1 μg/ml propidium iodide (PI) and 50 μg RNase and analyzed by flow cytometry (BD LSR-II) MCF-7 cells were exposed to 10 Gy γ radiation, were used as a positive control.

#### Double stranded DNA damage

Double standard DNA breaks were detected using p-H2AX (Ser139) antibody (Cell Signaling Technologies) using Immunofluorescence and western blotting method mentioned above.

### ROS measurement

ROS was measured using DCFDA dye (Thermo Fisher Scientific) using flow cytometry (BD LSR-II). Control and knockdown cells were washed with PBS and incubated with DCFDA dye (10 μM) for 30 minutes at 37°C. After incubation, cells were washed twice and suspended in 500 μL of PBS, then analyzed with flow cytometry. Data analysis was performed using FlowJo software. H202 was used as a positive control. Mitochondrial ROS was measured using MitoSOX-Red (Thermo Fisher Scientific) using flow cytometry (BD LSR-II). Control and knockdown cells were washed with PBS and incubated with MitoSOX-Red dye (10 μM) for 30 minutes at 37°C. After incubation, cells were washed twice and suspended in 500 μL of PBS, then analyzed with flow cytometry. Data analysis was performed using FlowJo software. Doxorubicin was used as a positive control.

### MTT assay

Cells were seeded at ~ 3000 – 10,000 cells/well in 96 well plates. After the cells were incubated overnight, they were treated with drugs. Drugs (obtained from Selleck chemicals or Sigma-Aldrich) were dosed from a clinically achievable low concentrations to clinically achievable high concentration, and four to six replicate wells were tested per concentration. Cells were treated with drugs for 48-120 hours. Thereafter, MTT dye (Thermo Fisher Scientific) was added to the cells for four hours, and SDS-HCl solution was added to each well and mixed thoroughly. After four hours of incubation, absorbance was measured at 570 nm. Cell survival is calculated as the percentage normalized to control.

## Declarations

### Consent for Publication

All authors have consented to this submission.

### Availability of data and material

GEO submission of microarray data is in progress.

## Acknowledgements

M.R.M. acknowledges the support of AAISCR. M.J.K and A.X.M. acknowledge the support of the graduate training programs in Cellular & Molecular Medicine and Biochemistry Cell and Molecular Biology, respectively, at the Johns Hopkins University. We would like to thank the Johns Hopkins Transcriptomics and Deep Sequencing Core for completing the microarray analysis.

## Author Contributions

M.R.M performed most experiments, including p53 signaling, ROS generation, DNA damage response and drug toxicity. M.J.K performed microscopy and image analysis. D.K.D performed cell cycle analysis and Ca^2+^ flux. J.W. and P.B. constructed and validated the TP53 mutant. A.X.M. and C.C.T analyzed the microarray data. M.R.M and R.R. wrote the paper, and all authors contributed to making the figures and editing the text.

